# HOCMO: A Tensor-based Higher-Order Correlation Model to Deconvolute Epigenetic Microenvironment in Breast Cancer

**DOI:** 10.1101/2020.12.01.406249

**Authors:** Charles Lu, Rintsen Sherpa, Liubou Klindziuk, Stefanie Kriel, Shamim Mollah

**Author notes:** These authors contributed equally to this work (Charles Lu, Rintsen Sherpa).

## Abstract

An in-depth understanding of epithelial breast cell response to growth-promoting ligands is required to elucidate how signals from the microenvironment affect cell-intrinsic regulatory networks and their resultant cellular phenotypes, such as cell growth, progression, and differentiation. Understanding the cellular response to these signals is particularly important in understanding the mechanisms of breast cancer initiation and progression. There is increasing evidence that aberrant epigenetic marks are present in cells of the breast tumor microenvironment and are known to affect these cellular processes. However, the mechanisms by which epigenetic microenvironment signals influence these cellular phenotypes are complex and currently not well established. To deconvolute the complexity of the epigenetic microenvironment signals in breast cancer, we developed a novel tensor-based correlation method: HOCMO (Higher-Order Correlation Model), applying to proteomics time series data to reveal the four-way regulatory dynamics among signaling proteins, histones, and growth-promoting ligands across multiple time points in the breast epithelial cells. HOCMO reveals two functional modules and the onset of specific protein-histone signatures in response to growth ligands contributing to distinct cellular phenotypes indicative of breast cancer initiation and progression. We evaluate robustness of our tensor model against baseline method TensorLy and achieved slight improvement in terms of reconstruction error and execution time. HOCMO is a data independent self-supervised learning method with superior interpretability that can capture the strength of complex interactions such as inter- and intra-pathway cellular signaling networks in any diseases or biological systems.

## Introduction

There is increasing evidence that epigenetic alterations in the cellular microenvironment affect normal tissue morphogenesis and promote breast cancer formation and progression^1^. The extracellular matrix (ECM), a major component of the cellular microenvironment, interacts with growth factors to influence cell behaviors contributing to growth, progression, and differentiation^2^. Details about these interactions within the complex and continuously changing breast microenvironment are important during tumor formation but are not well understood. Therefore, an in-depth understanding of these interactions within the tumor microenvironment is critical for the early detection and design of effective anti-cancer therapies. These complex interactions are controlled by higher-order inter- and intra-pathway cell signaling networks involving a cascade of biochemical reactions and physical interactions among proteins, genes, and other biomolecules. These interactions are typically represented as a network which encodes pairwise correlations between nodes with some mathematical formulation of their dynamics. Usual mathematical formulations of the dynamics typically assume that each node in the network has an associated “state” variable (representing, for example, expression of the corresponding signaling protein) as a function of the “time” variable. Describing the value of this state variable at a node (“state” of the node) depends on the state of other nodes interacting with it. A major drawback of this type of network is that it does not consider the higher-order correlations of the interactions, which may lead to an inaccurate or incomplete view of the global interconnectivity of the signaling proteins, and thus of their functional contributions to cellular phenotype. For example, a network only encodes pairwise correlations of node state variables and thus cannot represent a joint *k*-way correlation among *k* state variables for any *k* > 2. In a recent study, Mollah *et al*., developed a 3-way correlation network model, iPhDNet, to represent causal connectivity among phosphoproteins, histones, and drugs using a hybrid machine learning approach^3^. In this study, we develop a higher-order (k >= 3-way) correlation model using tensor decomposition from measured proteomic signals. In this study we develop a ligands-signaling proteins-histones-time 4-way correlation network by constructing a signaling proteins-ligands-time 3-way tensor from protein expression data and a histones-ligands-time 3-way tensor from histone modification data to understand the influences of various growth ligands on cell signaling pathways. We developed a Non-negative Tensor Factorization (NTF)^4^ based approach to capture the higher-order correlations among proteins, ligands and histones across multiple time points from the collected multi-omics data containing two time-series proteomic datasets. Unlike deep learning models, NTF learns the underlying complex representation of data in an unsupervised manner, offers better interpretability, scalability and data-efficiency when dataset is limited. Unlike deep learning models, NTF does not encounter the issue of being a black box model due to more transparent architecture. The overall workflow of our method consists of three components (Figure 1). First, we collect data from reverse phase protein array (RPPA) and global chromatin profiling (GCP) time series data and represent them as two 3D tensors. Then, we adopt an NTF based method to elicit the higher-order dynamic correlations encoded in the RPPA and GCP tensors. Finally, we focus on the interpretation of latent factors derived from the RPPA and GCP tensor factorization to collectively reveal the significant interactions among proteins, ligands, and histones at the respective time states. We introduce a novel metric called Higher-Order Correlation score (HOC-score) to quantify the correlation among ligands, phosphoproteins, and histones causing the chromatin structure changes.

**Figure 1.**
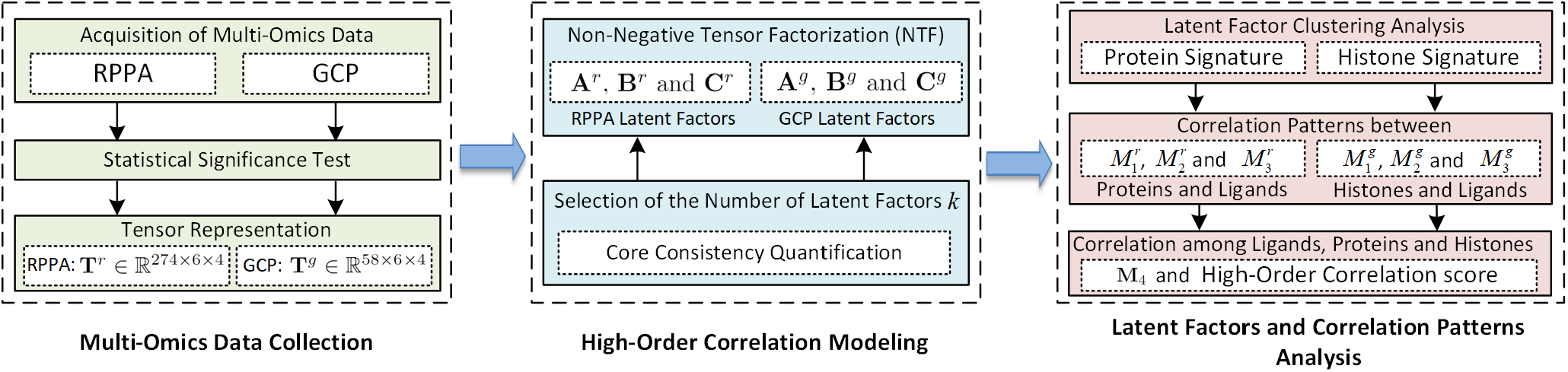
Workflow of the proposed approach. It involves three major components: i) multi-omics data preprocessing, ii) higher-order data correlation modeling using non-negative tensor factorization, and iii) interpretation of tensor factorization results.

A large-scale NIH initiative, the library of integrated network-based cellular signatures (LINCS) (http://www.lincsproject.org), has carried out the multi-omics characterization of breast epithelial cells to six growth-promoting ligands, through measurement of RPPA signaling proteins^5^, and GCP^6^. These measurements were carried out at multiple time points post treatment of cells. The RPPA and GCP are antibody and mass spectrometry (MS)-based targeted proteomics assays, respectively, where multiple growth ligands are used to treat a representative set of signaling proteins (RPPA) and different combinations of histone modifications (GCP). This higher-order tensor model offers a method for obtaining key signatures of cellular response to growth ligands based on detailed mechanisms reconstructed from these datasets. Specifically, we present an effective approach to elucidate how microenvironment signals affect cell-intrinsic intracellular protein signals and histone networks, leading to experimentally observable cellular phenotypes. Our methods identify histone modification fingerprints, which are endpoints of the complex signaling events following growth ligand treatment. Chromatin topology changes, caused by histone modifications reflect the altered cellular state.

For characterization of the breast microenvironment responses, we used the human breast epithelial cell line (MCF10A) from the LINCS study which profiled 295 signaling proteins and 65 histone marks at 4, 8, 24, and 48 hours after treatment with six growth-promoting ligands: epidermal growth factor (EGF), hepatocyte growth factor (HGF), oncostatin M (OSM), bone morphogenetic protein 2 (BMP2), transforming growth factor beta (TGFB), and interferon gamma-1b (IFNG). These ligands target diverse receptor types and activate various canonical downstream pathways and were chosen because they elicited multiple phenotype changes affecting proliferation, migration, and morphology. To evaluate our results from proteomics analyses, we further performed differential expression analysis and gene set enrichment analysis to capture gene activities at the transcriptomic level using RNAseq data.

## STAR Methods

### Multi-Omics Datasets

We utilized the RPPA and GCP datasets from LINCS to capture the dynamic regulations of growth ligands influencing phosphoproteins and histone modifications in the breast tumor microenvironment. We then used the RNAseq dataset from LINCS to further verify our phosphoprotein-histone signatures results with the respective transcriptional profiles.

### RPPA and GCP Data Acquisition

We downloaded the level3 (log 2 normalized) RPPA data from the Synapse platform (https://www.synapse.org/#!Synapse:syn12555331) in June 2020. The dataset consists of highly sensitive and selective antibody-based measurements of 295 proteins and phosphoproteins after individual treatments with six growth promoting ligands, namely, EGF, HGF, OSM, BMP2, TGFB, and IFNG. Each data sample contains three replicates for each of these ligands and a control treated with phosphate-buffered saline (PBS) measured at five time points (1, 4, 8, 24, and 48 hours).

We downloaded the level 3 (log2 normalized) GCP data from the Synapse platform (https://www.synapse.org/#!Synapse:syn18804314) in August 2020. This dataset consists of 65 probes monitoring combinations of post-translational modification on histones in MCF10A healthy breast epithelial cell line. Cells in these assays were treated with EGF, HGF, OSM, BMP2, TGFB, or IFNG at various concentrations. The data contains three replicates for each of these ligands and a control treated with PBS. These samples were measured at four time points (4, 8, 24, and 48 hours).

To harmonize data and provide analyses that are consistent across the above two datasets, we removed all samples in the first hour from the RPPA data. This resulted in RPPA and GCP data consisting uniform four distinct time points of 4, 8, 24 and 48 hours. In both RPPA and GCP datasets, each data sample (e.g., protein or histone mark) has three replicates representing three independent experiments at each time point. We applied an unpaired Student’s *t*-test to retain statistically significant (*p*-value<0.05) 274 proteins in RPPA and 58 histones in GCP in the experiment.

### RNAseq Data Acquisition

The level 0 (fastq) RNAseq data were downloaded from the Synapse platform (https://www.synapse.org/#!Synapse:syn18518040) in January 2021. 100bp single-end reads were sequenced on an Illumina HiSeq 2500 Sequencer by OHSU MPSSR. The dataset contains the gene expression data of MCF10A breast cells treated with six growth ligands: EGF, HGF, OSM, BMP2, TGFB, and IFNG. Each data sample contains three replicates for each ligand measured at 24 and 48 hours (EGF has four replicates per timepoint). Cells were also treated with a control (PBS) and measurements were taken at 0, 24, and 48 hours. The fastq files were aligned to the ENSEMBL GRCh38 assembly of the human genome using the STAR sequence aligner^8^ version 2.7.3 and output as BAM files. A count matrix of all samples and their genes was generated using the featureCounts program^9^ from the R package RSubread^10^ version 2.6.1 and the ENSEMBL GRCh38 gene annotations.

### Tensor Representation of RPPA and GCP Data

After performing the significant test and sample screening processes, the differential abundance (fold change, *i.e*., the ratio of each sample measurement to its control measurement) of signaling proteins and histones relative to their respective controls (PBS) are calculated. Figure 2A illustrates the differential abundance of an example protein PIK3CA from RPPA data and an example histone mark H3K9me2S10ph1K14ac0 from the GCP data. For example, after treatment with IFNG, expression of the protein PIK3CA was initially decreased during the 4^*th*^ and 8^*th*^ hours and was then elevated until the 48^*th*^ hour. Meanwhile, expression of the histone mark H3K9me2S10ph1K14ac0 showed consistent elevation from the 4^*th*^ to 48^*th*^ hour.

**Figure 2.**
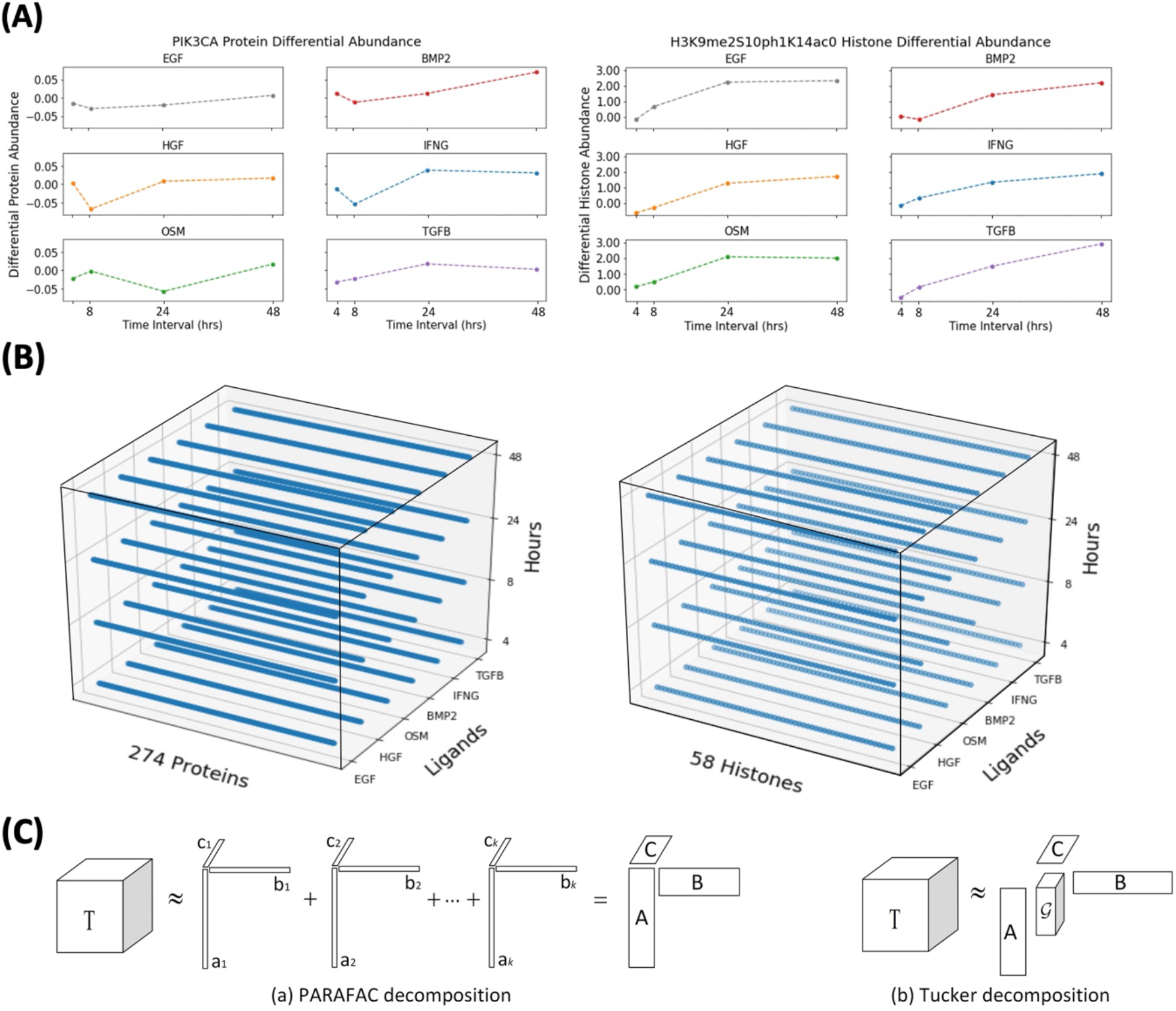
(A) Differential abundance changing over time. The differential abundances of protein PIK3CA and histone H3K9me2S10ph1K14ac0 after treatment with six growth ligands at various time points. **(B) Visualizations of the 3-way RPPA tensor and GCP tensor**. Each tensor has three dimensions or modes, protein, ligand and time (hour) for RPPA (left), and histone, ligand and time (hour) for GCP (right). **(C) The graphical representations of PARAFAC and Tucker**. Tucker contains a core tensor *𝒢* and PARAFAC is a special case of Tucker when the core tensor *𝒢* is super-diagonal.

The RPPA fold-change data at each time point *i* with 274 proteins and 6 ligands can be represented in a matrix 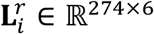, where the superscript *r* denotes RPPA data. By stacking all matrices observed at four sequential time points (e.g., 4, 8, 24 and 48 hours), we are able to represent the RPPA data with a 3D tensor (a.k.a 3-way array) denoted by 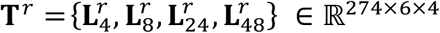. Similarly, The GCP fold-change data with 58 histones and 6 ligands can be represented as a 3D tensor 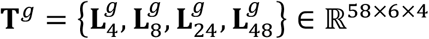 (with superscript *g* denoting GCP data), where 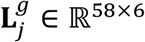 denotes the fold-change matrix at time point *j*. **T**^*r*^ and **T**^*g*^ are visualized in two 3-way tensors shown in Figure 2B (left) & (right) respectively.

The differential abundance analysis on RPPA and GCP data sets captures fold-change values reflecting negative and positive regulations of phosphoproteins and histone abundances incurred by the treatment of the six growth ligands. To satisfy the non-negativity constraint of the NTF method, we first transformed these fold-change values to their absolute values. Using these absolute fold-change values we then construct two tensors **T**^*r*^ **= T**^274×6×4^ for RPPA representing fold-changes of 274 phosphoproteins and **T**^*g*^ **= T**^58×6×4^ for GCP representing fold-changes of 58 histone modifications after the treatment of six growth ligands across four (4-, 8-, 24-, and 48-hour) time points (Figure 2B). We used these tensors as inputs for the subsequent non-negative tensor factorization steps.

### Higher-Order Correlation Modeling

The tensors encode for the correlations or interactions between proteins and ligands (or between histones and ligands) changing over time (Figure 2B). In this section, we present a framework using NTF to model multi-layer correlations (4-way correlations), such as those among ligands, proteins, histones, and time shown above.

#### Non-Negative Tensor Factorization (NTF)

Tensor factorization can be viewed as the higher-order version of matrix factorization^4^ by extending the two-dimensional matrices to tensors. A tensor is a multi-way array or multi-dimensional matrix^11^ formally defined as:

**Definition 1 (Tensor)** *Let I*_1_, *I*_2_, ⋯, *I*_*N*_ *denote index upper bound. A tensor* 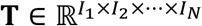 *of order N is a N-way array where elements* 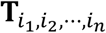 *are indexed by i*_*N*_ ∈ {1,2, ⋯, *I*_*N*_} *for* 1 ≤ *n* ≤ *N*.

Tensors are generalizations of vectors and matrices, where the rank of a tensor is the minimum number of rank-one tensors defined as:

**Definition 2 (Rank-One Tensor)** *A tensor* 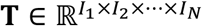 *of order N has rank-one if it can be written as the outer product (represented by notation* ∘*) of N vectors, i*.*e*., **T = a**^(1)^ ∘ **a**^(2)^ ∘ ⋯ ∘ **a**^(*N*)^ *or* 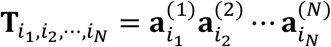 *for all* 1 ≤ *i*_*N*_ ≤ *I*_*N*_.

Tensors can capture the complex relations between a larger number of factors, such as the interplay between protein, ligand and time factors in RPPA data, with higher fidelity. In turn, tensor factorization provides a way to decompose a given tensor into *k* components or latent factors in different domains with common latent factors reflecting the correlations across these different domains. A model which imposes non-negativity on all resulting components is called NTF. The two widely used tensor factorization techniques are the Parallel Factor Analysis (PARAFAC)^12^ and Tucker^13^, which have shown promises in many real-world applications such as cancer survival prediction^14^ and image processing^15^.

#### PARAFAC and Tucker

The non-negative PARAFAC was proposed by Carroll *et al*.^16^ and then developed by Bro^17^ with a more efficient method to calculate the non-negative components. Figure 2C (a) shows the graphical representation of PARAFAC, which aims to decompose a 3-way tensor 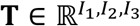 into a sum of component rank-one tensors as:

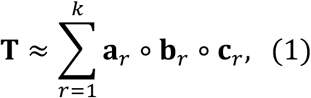

where 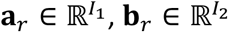 and 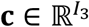. *k* represents the number of latent components or factors approximating the actual rank of tensor **T**. Data relationships in the original tensor (**T**) are thereby fully preserved by the resulting components **a**_r_, **b**_r_, and **c**_r_. Alternatively, the PARAFAC decomposition can be represented as:

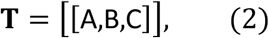

where the loading matrices **A**,**B**,**C** can be understood as the linear mapping from the respective modes to the latent spaces defined as:

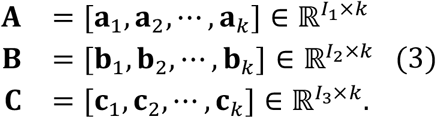

Tucker decomposition is another popular NTF technique^13^. As shown in Figure 2C (b), compared with PARAFAC, in Tucker an additional core tensor 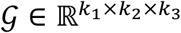 is calculated together with the three loading matrices **A, B** and **C** represented as:

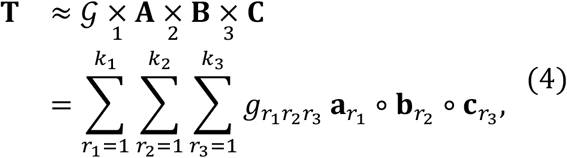

where *k*_1_ × *k*_2_ × *k*_3_ rank-one tensors are produced, unlike PARAFAC which only has *k* rank-one tensors. *k*_1_, *k*_2_ and *k*_3_ are numbers of components for each mode of the 3-way tensor.

Therefore, PARAFAC is a special case of Tucker with the super-diagonal core tensor *𝒢*. Studies^18,19^ show that PARAFAC is suitable for problems where the latent factor interpretation is essential, whereas Tucker is often used for data compression purpose since the core tensor can be viewed as a compression of the original tensor. Another difference is that the rank-one decomposition tensors of PARAFAC are often unique, whereas Tucker decomposition is not^20^. For these reasons, we chose to use non-negative PARAFAC to analyze and study the tumor microenvironment. There are many methods to implement the 3-way PARAFAC including Nonnegative CANDECOMP/PARAFAC (NCP)^21^ and TensorLy^22^. In this study we chose to implement NTF using NCP for our RPPA and GCP tensors given its lower reconstruction error as shown in Figure 3A, and the importance of interpretability in a biological context. The NTF package as part of TensorLy was used to validate our results obtained from NCP.

**Figure 3.**
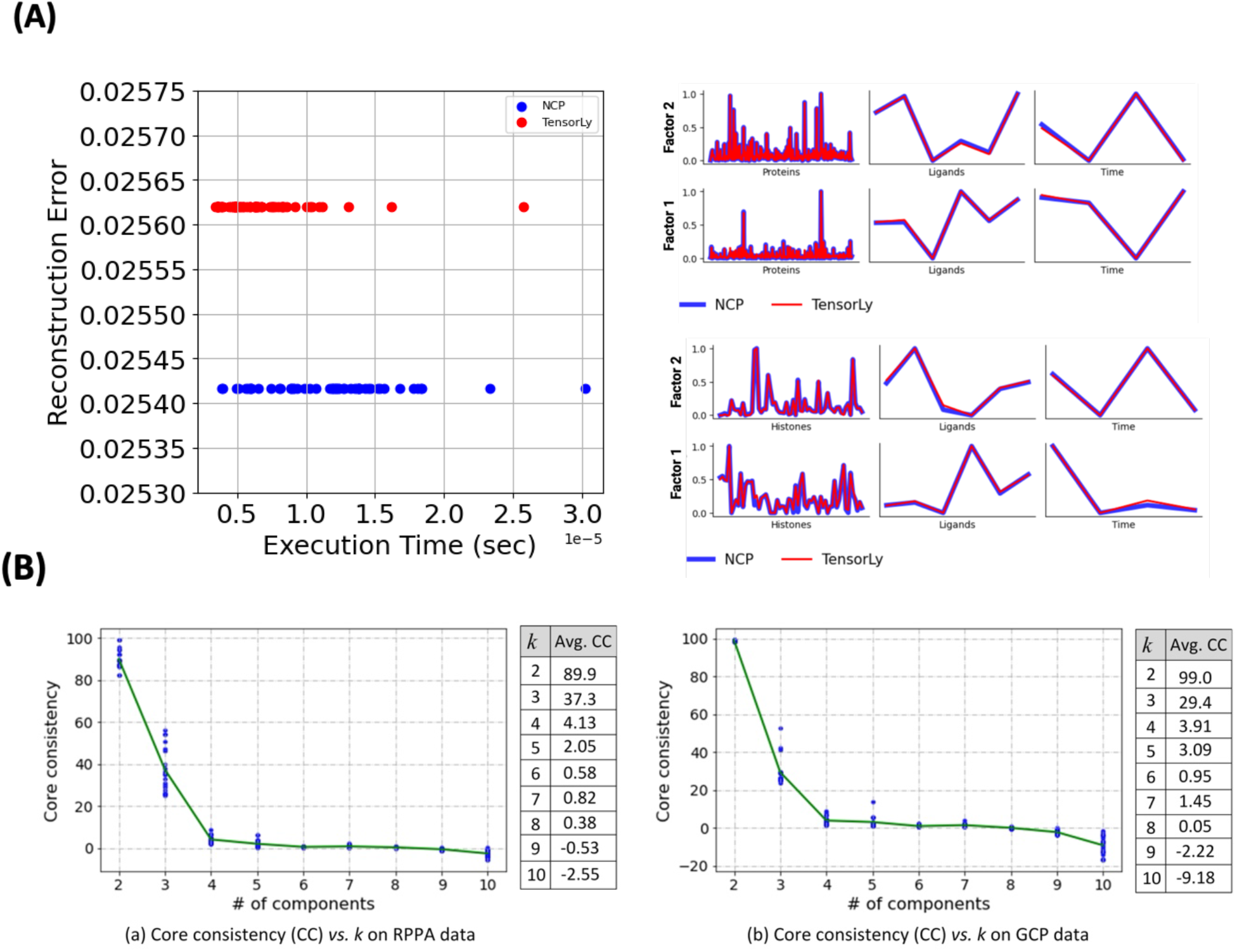
(A) *Evaluation* of *the* model. Factors derived from the tensor decomposition processes (TensorLy and NCP) were used to reconstruct a tensor. The mean squared error was measured between this reconstructed tensor and our original tensor and plotted against the execution time of the tensor decomposition process (left). Comparison of the performance of NCP and TensorLy implementations using two factors to estimate protein, histone, ligands and time in RPPA and GCP data(right). **(B) The core-consistency vs. the number of components (***k*). The core consistency for each *k* tested on the RPPA tensor (left) and GCP tensor (right) was computed 100 times independently. The top 20% core consistencies (blue dots) with their averages (solid line) are shown in the left panel (graph). The actual values for average core consistencies (CC) are shown in the right panel (table).

#### Non-Negative PARAFAC Based on NCP

NCP has been widely used to solve the non-negative PARAFAC decomposition, which is implemented based on the Block Principal Pivoting (BPP) method^23^. For a given 3-way tensor 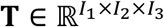, NCP aims to minimize the following objective:

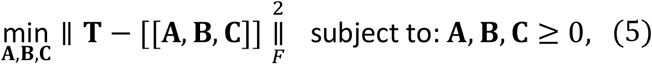

In other words, NCP aims to solve latent factor matrices 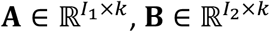 and 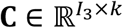 (*k* is the number of latent factors) to approximate the input tensor **T**. In the following, we briefly describe the principle of solving **A, B** and **C** in NCP. For mathematical simplicity, we use **A**^(1)^, **A**^(2)^ and **A**^(3)^ to respectively denote **A, B** and **C** in the paragraph. For the GCP method, the tensor approximation **T** ≈ [[**A, B, C**]] can be reformulated as:

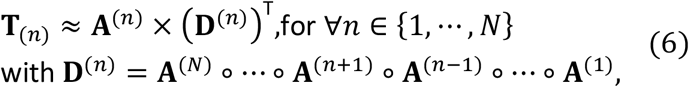

where *N* **=** 3 representing the 3-way tensor factorization in this paper and **T**_(*N*)_ represents the mode-*n* matricization of tensor **T**. Then, the optimization in Eq.5 can be viewed as solving the following subproblems:

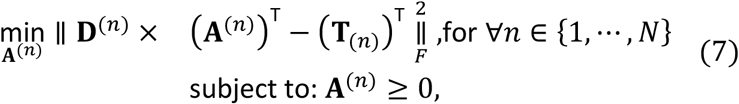

where we need to solve **A**^(1)^, **A**^(2)^ and **A**^(3)^ in this paper. From Eq.7 we can see that NCP actually reduces the NTF to *N* independent non-negative matrix factorization problems, where each problem can be simplified as:

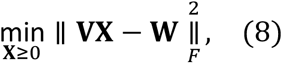

where **V** ∈ ℝ^*p*×*Q*^ corresponds to **D**^(*N*)^, **X** ∈ ℝ^*Q*×*L*^ corresponds to (**A**^(*N*)^)^T^, and **W** ∈ ℝ^*p*×*L*^ corresponds to *N*(**T**^(*N*)^)^T^ where *P, Q, L* are integers. Then, the BPP algorithm with multiple right-hand sides is utilized to solve these subproblems.

### Selection of the Number of Latent Factors *k*

Before applying the non-negative PARAFAC on the multi-omics data, the number of latent components or factors *k* needs to be determined to approximate the actual rank of the target tensor.

It is non-trivial to determine the best number of latent factors *k*, which typically necessitates multiple trials and fine-tuning^24^. The *core consistency* (CC) metric has been widely used to evaluate the fitness of a candidate *k*,^25^ where the resulting latent components of an appropriate PARAFAC model are reflective of the trilinear variation of the data. Since the Tucker decomposition can represent the trilinear variation in data by the shared core tensor and various loading matrices, the definition of CC is naturally achieved by examining the difference between PARAFAC and Tucker based tensor decomposition. As demonstrated in the previous section, the PARAFAC decomposition is a special case of the Tucker decomposition (Eq.4) since Eq.1 can be rewritten as:

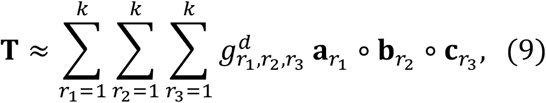

where 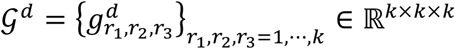 is a super-diagonal tensor with all diagonal elements (e.g., *r*_1_ **=** *r*_2_ **=** *r*_3_) equal to 1 and other off-diagonal elements equal to 0. The Tucker core tensor *𝒢* can be considered as the optimal representation of the target tensor **T** w.r.t. the subspace defined by the latent factors^25^. In this regard, if a specific *k* is optimal for PARAFAC to fully span trilinear variations in the data, the Tucker core tensor *𝒢*, computed with fixed loading matrices output from PARAFAC upon *k*, should be close to the super-diagonal. Then, the CC is defined as the difference between *𝒢* and *𝒢*_*d*_ to quantify the goodness of a particular *k*:

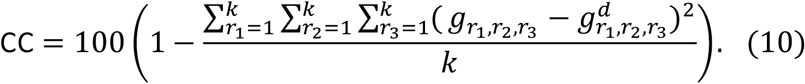

The CC reflects how well a particular Tucker core fits the assumption of a good PARAFAC model for a given *k*. In order to accurately represent the data, typically a larger value of *k* is chosen such that a high CC value is yielded (max value = 100). A CC above 50 is interpreted as an acceptable model^24,25^.

Instead of choosing the number of latent factors randomly, we leverage CC as the metric to determine the optimal number of latent factors *k* for RPPA and GCP tensors. Using Eq.10, we calculated CC values for *k* ranging from 2 to 10 (Figure 3B). We observed the best average core consistencies for RPPA and GCP tensors at 89.9 and 99.0 respectively when *k* **=** 2. Although a larger *k* would be desirable, the CC values for all *k* > 3 for both RPPA and GCP data are less than 50 (a value that is considered as a not acceptable model). Therefore, in this study, we consider *k* **=** 2 as the optimal number of latent components for PARAFAC decomposition on both RPPA and GCP data.

### NTF on RPPA and GCP Tensors

After determining the optimal number of components by core consistency, we applied the non-negative PARAFAC model implemented in NCP to decompose the RPPA tensor **T**^*r*^ ∈ ℝ^274×6×4^ and GCP tensor **T**^*g*^ ∈ ℝ^58×6×4^, respectively. The non-negative factorization on RPPA data based on NCP is summarized in Algorithm 1, which takes the 3-way RPPA tensor **T**^*r*^ as an input and then decomposes it to three latent factor matrices **A**^*r*^, **B**^*r*^ and **C**^*r*^ (*r* represents RPPA tensor). Nonnegative Double Singular Value Decomposition (NNDSVD)^26^ is applied to initialize **A**^*r*^, **B**^*r*^ and **C**^*r*^, based on two Singular Value Decomposition (SVD) processes, with one approximating the data matrix and the other approximating positive sections of the resulting partial SVD factors utilizing an algebraic property of unit rank matrices. Then the block principle pivoting (BPP) is applied to generate the final loading (latent factor) matrices **A**^*r*^, **B**^*r*^ and **C**^*r*^ (Algorithm 1).

#### Algorithm 1

RPPA tensor decomposition based on NCP

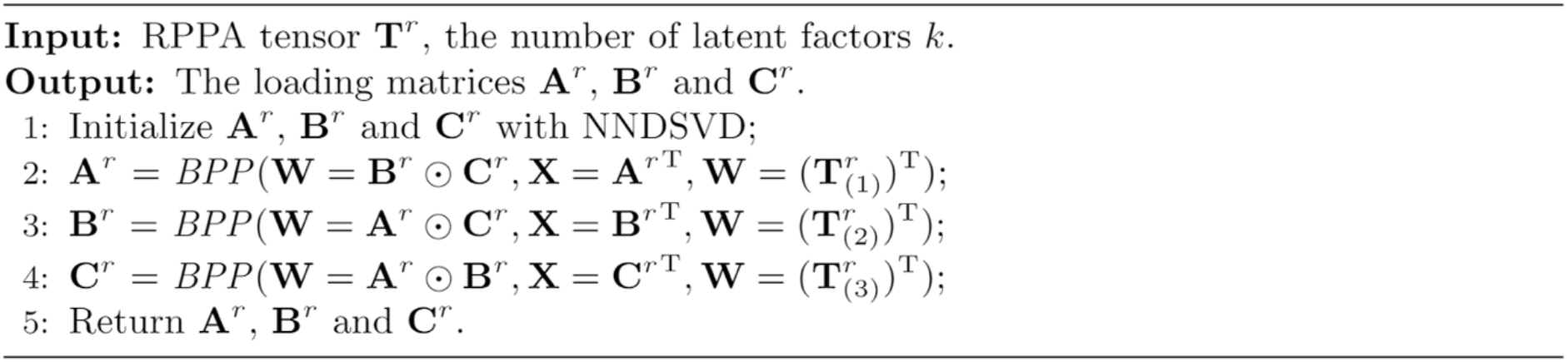

The same algorithm is applied to decompose the GCP tensor **T**_*g*_ and derive the latent factor matrices **A**^*g*^, **B**^*g*^ and **C**^*g*^ (*g* represents GCP tensor).

### Higher-Order Correlations Representation

To gain insights into the interactions between proteins and ligands in the RPPA data over time, we further constructed three auxiliary matrices through the combinations between 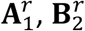 and 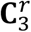 as:

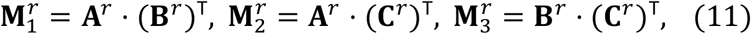

where ⋅ denotes the matrix multiplication. 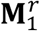 (Supplemental Table 1) stores the pairwise interaction intensities between 274 proteins and 6 ligands, while 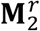 (Supplemental Table 2) and 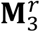 (Supplemental Table 3) store the pairwise interaction intensities of 4 time states with the 274 proteins and 6 ligands respectively. To reveal the dynamic correlations between histones and ligands in the GCP data, we created three matrices as follows:

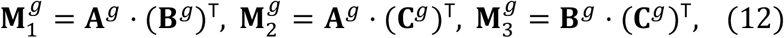

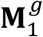 (Supplemental Table 4) captures the pairwise interaction intensities between the 58 histones and 6 ligands, while 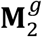 (Supplemental Table 5) and 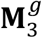 (Supplemental Table 6) store the pairwise interaction intensities of 6 time states with 58 proteins and 6 ligands respectively. Further, based on the two matrices 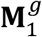 and 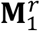, *i*.*e*., we constructed **M**_4_ (Supplemental Table 7) that correlates histones and proteins from the RPPA and GCP data to represent their pairwise correlations by:

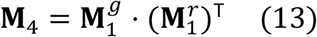

In this study, we use **M**_4_ (e.g., positive values) to quantify the relative interaction or correlation intensity between proteins and histones after the treatment of six growth ligands. The directions (e.g., either positive or negative) of protein and histone regulations incurred by the growth ligands at respective time points can be obtained from the indicator tensors *I*^*r*^ **=** {−1,1}^274×6×4^ for RPPA and *I*^*g*^ **=** {−1,1}^58×6×4^ for GCP. To better understand the epigenetic tumor environment of breast tissue, we aim to elucidate the regulatory mechanisms governing histone modifications mediated by ligand-induced phosphoproteins and their consequential impact on changes in chromatin structure. To quantify the correlation amongst ligands, phosphoproteins, and modifying histones, we define a Higher-Order Correlation (HOC) score for any triplet relation <i, j, k> representing the i^th^ ligand, j^th^ phosphoprotein and kth histone as:

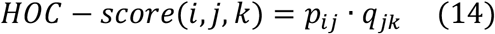

where *p*_*ij*_ is the interaction intensity between the ith ligand and jth protein stored in 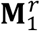, and *q*_*ik*_ is the interaction intensity between the jth protein and kth histone stored in **M**_4_. As the value of HOC approaches 1, the strength of correlation among triplets at a specific time point increases.

### Gene Set Enrichment Analysis

For validation, we performed a gene set enrichment analysis to find out which canonical pathways associated with the significant phosphoproteins identified from NTF were most affected by each ligand treatment. Using the RNAseq raw count matrix data, an expression dataset file and a phenotype label file were created as specified in the GSEA User Guide for each individual ligand at a specific time point compared to PBS at the same time point. GSEA^27^ version 4.1.0 was run for each comparison using ‘c2.cp.kegg.v7.2.symbols.gmt’ as the gene set database, ‘Human_ENSEMBL_Gene_ID_MSigDB.v7.4’ as the chip platform, and ‘phenotype’ as the permutation type, set at 1000 permutations.

## Results

### HOCMO captures histone signatures of tumor microenvironment (TME) structures

*Latent Factor Analysis*: In order to find patterns within the protein and histone modification datasets, we decomposed the RPPA and GCP tensors into their latent factors. These latent factors were mapped onto a shared two-dimensional space and represented by matrices for each variable (Figure 4A). We observed variable affinities towards the two latent components within each of the matrices. Distinct groups can be easily seen. For both RPPA (Figure 4A a ***C***^*r*^) and GCP (Figure 4A b ***C***^*r*^) datasets, the 4, 8, and 24 hour time states appear to map closely to the second latent factor, while the 48hr time state has a greater affinity for the first latent factor. In the histone latent matrix (Figure 4A b ***A***^*g*^), two groups of histones can be seen separated by the latent factors. However, not all affinities are immediately clear from the latent matrices, and in order to properly define groups of related proteins and histones, we must further perform clustering analysis.

**Figure 4.**
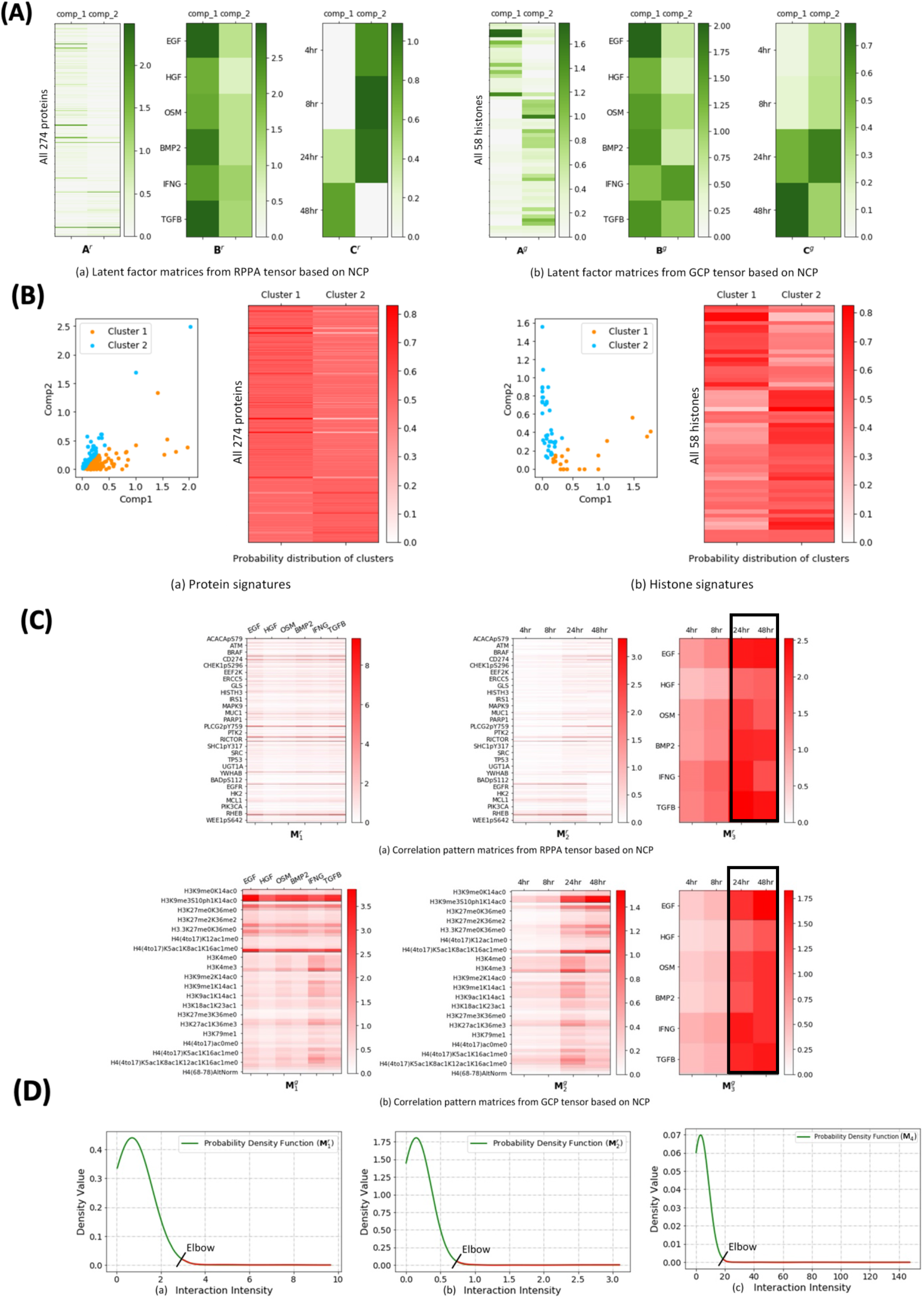
(A) Latent factors generated from NTF. (left) *A*^*r*^, *B*^*r*^ and *C*^*r*^ represent proteins, ligands and hours from RPPA that are mapped to a two-dimensional latent space. Rows in *A*^*r*^, *B*^*r*^ and *C*^*r*^ represent all 274 proteins, 5 ligands and 4 time states from the RPPA data respectively, and columns represent the two shared latent factors obtained by NTF. (right) *A*^*g*^, *B*^*g*^ and *C*^*g*^ represent histones, ligands and hours from GCP that are mapped to a two-dimensional latent space. Rows in *A*^*g*^, *B*^*g*^ and *C*^*g*^ represent all 58 histones, 5 ligands and 4 time states from the GCP data respectively, and columns represent the two shared latent factors obtained by NTF. **(B) The identified (left) protein signatures and (right) histone signatures**. We performed clustering analysis based on the factor matrices ***A***^*r*^ *and* ***A***^*g*^ respectively. In the scatter plots, each cluster corresponds to a latent component and each protein or histone belongs to the cluster with largest membership value. The heatmaps in (left) and (right) show the actual probability distributions of each protein and histone to the respective clusters, where rows contain 274 proteins (only 28 protein labels shown) and 58 histones (only 20 histone labels shown) respectively. The protein signatures can be interpreted as the categorization of proteins demonstrating similar responses after treated with six growth ligands. Similarly, histones in the same cluster demonstrate similar responses after treated with these same growth ligands. **(C) The generated pattern matrices from the latent factor matrices**. Rows in 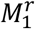 and 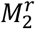 represent all 274 proteins, while their columns represent 6 ligands and 4 time states respectively (top panel). 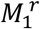 reveals the correlation patterns between 274 proteins and six ligands, where the onsets are revealed in 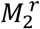 and 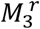 over the 4 time points in the RPPA data. (bottom) Rows in 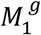 reveals the correlation patterns between 58 histones and 6 ligands, where the onsets are revealed in 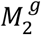 and 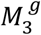 over the 4 time points in the GCP data. **(D) The distribution of intensity values for correlation pattern matrices** 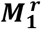 **and** 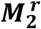. A density distribution learning is performed on (a) 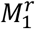 between signaling proteins and growth ligands, (b) 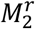 between signaling proteins and time points, and (c) 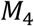 between signaling proteins and histones. Correlations with interaction intensities above the elbow are considered to be significant.

*Distinct clusters for phosphoproteins and histone modification signatures through latent structure:* To visualize related groups of proteins and histones based on their latent factors, we clustered the protein latent matrix **A**^*r*^ (Figure 4A a) and the histone latent matrix **A**^*g*^ (Figure 4A b). Each protein/histone was assigned to a cluster based on which of the two latent factors it had a greater affinity towards, resulting in two clusters within proteins (Figure 4B a) and histones (Figure 4B b). In the tumor microenvironment, the protein signatures can be interpreted as the categorization of proteins demonstrating similar responses to the growth ligands. Similarly, histones in the same signature may correspond to co-occurring histone modifications in response to the six growth ligands. In addition to the above hard clustering, we applied a Softmax transformation, visualized as a heatmap, which reveals the probability distributions of different proteins/histones belonging to each cluster.

### Correlation patterns between proteins and ligands and histones and ligands reveal potential onset at 24th hour

To further quantify the interactions between proteins and ligands, as well as the variations of their interactions over time, we constructed pairwise interaction matrices for each of the three pairs between proteins, ligands, and time states (Figure 4C top panel, Supplemental Table 8) based on their latent factor matrices (Figure 4A a). We can see that proteins are variably regulated by the six growth ligands (Figure 4C top panel 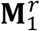), *i*.*e*., if a high intensity value (e.g., the value is larger than a selected threshold) is observed between a protein and a ligand in the matrix, we can understand that the protein has been significantly regulated by the corresponding ligand. Additionally, we see that proteins are regulated by ligands at different time states, where most proteins and ligands are significantly active at the 24-hour time point (Figure 4C top panel 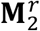 and 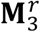). Similarly, we constructed pairwise interaction matrices for each of the three pairs between histones, ligands, and time states (Figure 4C bottom panel, Supplemental Table 9) based on their latent factor matrices (Figure 4A a). We can see that some histones are significantly regulated by the respective ligands (Figure 4C bottom panel 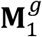). We also see parallels to correlations found in the protein data, as histones and ligands appear to be significantly active after the 24-hour time point (Figure 4C bottom panel 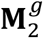 and 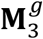.

### Identify significant interactome of ligands, signaling proteins, and histones triplets

To gain insights into the mechanism of how ligands regulate proteins and further mediate histones, we obtained all the significant proteins and histones based on their interactions with ligands (Figure 4C top panel 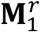 and Figure 4C bottom panel 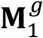). We are interested in investigating the most significant interactions and the respective driver proteins, histones, and ligands. First, to identify significantly interacting proteins and histones, we performed a density distribution learning (e.g., fitting a Gaussian distribution, Figure 4D (a)) over the intensity values in the interaction matrix between proteins and histones (Figure 4C top panel 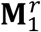). Note that since all the original values in the correlation matrices are non-negative, the fitted curve presents as a truncated Gaussian distribution displaying only the non-negative portion. We chose a cut-off value at the elbow as the threshold to determine significance, which has been a widely used method to decide the threshold in existing works^28,29^. If the interaction intensity between a protein and a corresponding ligand is larger than the threshold, we then consider that the protein has been regulated by the respective ligand. By considering the above two factors, we finally filtered out eight significantly regulated proteins and phosphoproteins: 1) CCNB1, CDC2, PLK1, RB1pS807S811, RPS6pS240S244, DUSP4 and RPS6pS235S236 were regulated by all six ligands, and 2) MYH2 was regulated by EGF, BMP2 and TGFB. We then perform a similar density distribution learning (Figure 4D (b)) on the interaction matrix between proteins and time states (Figure 4C top panel 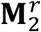) and found that the above eight phosphoproteins are all significant after the 24-hour time point.

Similarly, to determine significantly active histone marks in response to the growth ligands, we once again performed density distribution learning, this time over all intensity values in the interaction matrix between histones and ligands (Figure 4C bottom panel 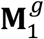), as well as the interaction matrix between histones and time states (Figure 4C bottom panel 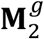). We finally filtered out the following five significant histone marks: H3K9me2S10ph1K14ac0, H3K9me3S10ph1K14ac0, H3K18ub1K23ac0, H4(20to23)K20me0 and H3K4ac1. Histones H3K9me2S10ph1K14ac0, H3K9me3S10ph1K14ac0 and H4(20to23)K20me0 were regulated by all six growth ligands, while histone H3K18ub1K23ac0 was regulated by EGF and TGFB, and histone H3K4ac1 was specifically regulated by IFNG. All histones were significantly active in the 24th and 48th hours observed except for H3K4ac1 which was active only in the 24th hour. Among the selected histones, H3K9me2S10ph1K14ac0, H3K9me3S10ph1K14ac0 and H3K18ub1K23ac0 are repressive histone marks, while H4(20to23)K20me0 is an active histone mark.

It is worth noting that all six ligands were significantly active at the 24th and 48th hour time points based on the intensity distributions on the respective interaction matrices (Figure 4C top panel 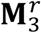 and Figure 4C bottom panel 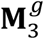). Then, we further explored the correlations between histones and proteins to identify which of the eight identified proteins contributed to the modifications of the five identified histones listed above. Based on the interactions between proteins and ligands (Figure 4C top panel 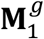) together with the interactions between histones and ligands (Figure 4C bottom panel 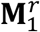), we constructed an additional matrix (**M**_4_: Figure 5a) to represent the pairwise interactions between all 274 proteins and 58 histones. We then performed a Gaussian distribution fitting the density distribution of the interaction intensities, where all interactions with coefficients over the cut-off value at the elbow on the curve were considered as significant (Figure 4D (c)). This resulted in eight significant proteins that have contributed to the modification of the five identified significant histones. In order to verify the above correlations, we also performed NCP and TensorLy based tensor factorization, which have produced the same results.

**Figure 5.**
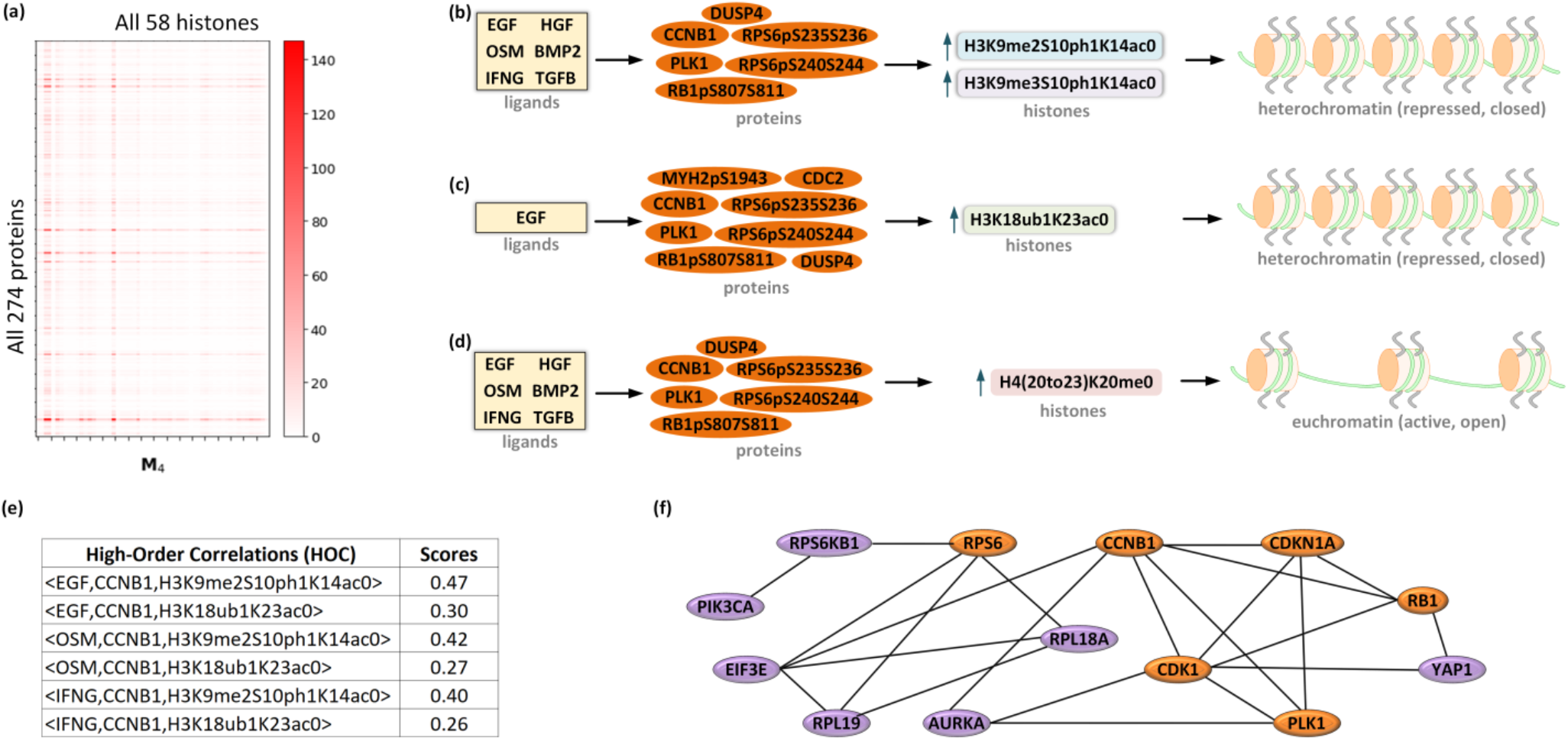
Significant signaling proteins and histones interactions. (a) Showing the HOC-based interactions between signaling proteins and histones. (b), (c) and (d) Showing specific ligands mediated protein and histone regulations, where four significant histones and eight proteins are examined. (e) Showing examples of the higher-order correlation (HOC) scores for six triplet relations among ligands, proteins and histones. (f) Showing the protein-protein interaction network among six enriched proteins (orange, obtained from HOCMO) and seven inferred proteins (purple) obtained from STRINGdb.

Finally, we summarized the identified correlations between proteins and histones after the treatment of growth ligands in three groups as hypothetical pathways: 1) Six proteins (CCNB1, PLK1, RB1pS807S811, RPS6pS240S244, DUSP4 and RPS6pS235S236) were significantly regulated after treatment with the six ligands, which then led to the modifications of the two repressive histone marks H3K9me2S10ph1K14ac0 and H3K9me3S10ph1K14ac0 resulting a repressed chromatin state (e.g., closed status of chromatin) (Figure 5b); 2) The growth ligand EGF regulated all eight proteins, which then mediated the active histone mark H3K18ub1K23ac0 which led to a closed chromatin conformation (Figure 5c); 3) Six growth ligands regulated six proteins (CCNB1, PLK1, RB1pS807S811, RPS6pS240S244, DUSP4 and RPS6pS235S236) which mediated histone mark H4(20to23)K20me0 resulting in an open chromatin conformation (Figure 5d). In order to further quantify these chain-based regulation, we developed a metric called Higher-Order Correlation score (HOC-score) which considered the correlation among ligands, proteins, and histones at a given time point. HOC-score reveals how much the respective interaction among ligands, proteins and histones contributes to the chromatin structure changes (Figure 5e). We further inferred interactive proteins of the identified seven enriched proteins using STRINGdb^30^ (Figure 5f).

### Identify signaling networks that controls cellular phenotypes: mapping to canonical signaling pathway

In order to contextualize these findings, we further mapped the eight enriched proteins and phosphoproteins to their corresponding canonical breast cancer signaling pathways using the KEGG and Reactome databases^31^ (Figure 6). We observed strong upregulation of all these eight proteins and phosphoproteins across all ligands after the 24-hour mark. Differential expression within these canonical signaling pathways infers disruption of healthy cellular function.

**Figure 6.**
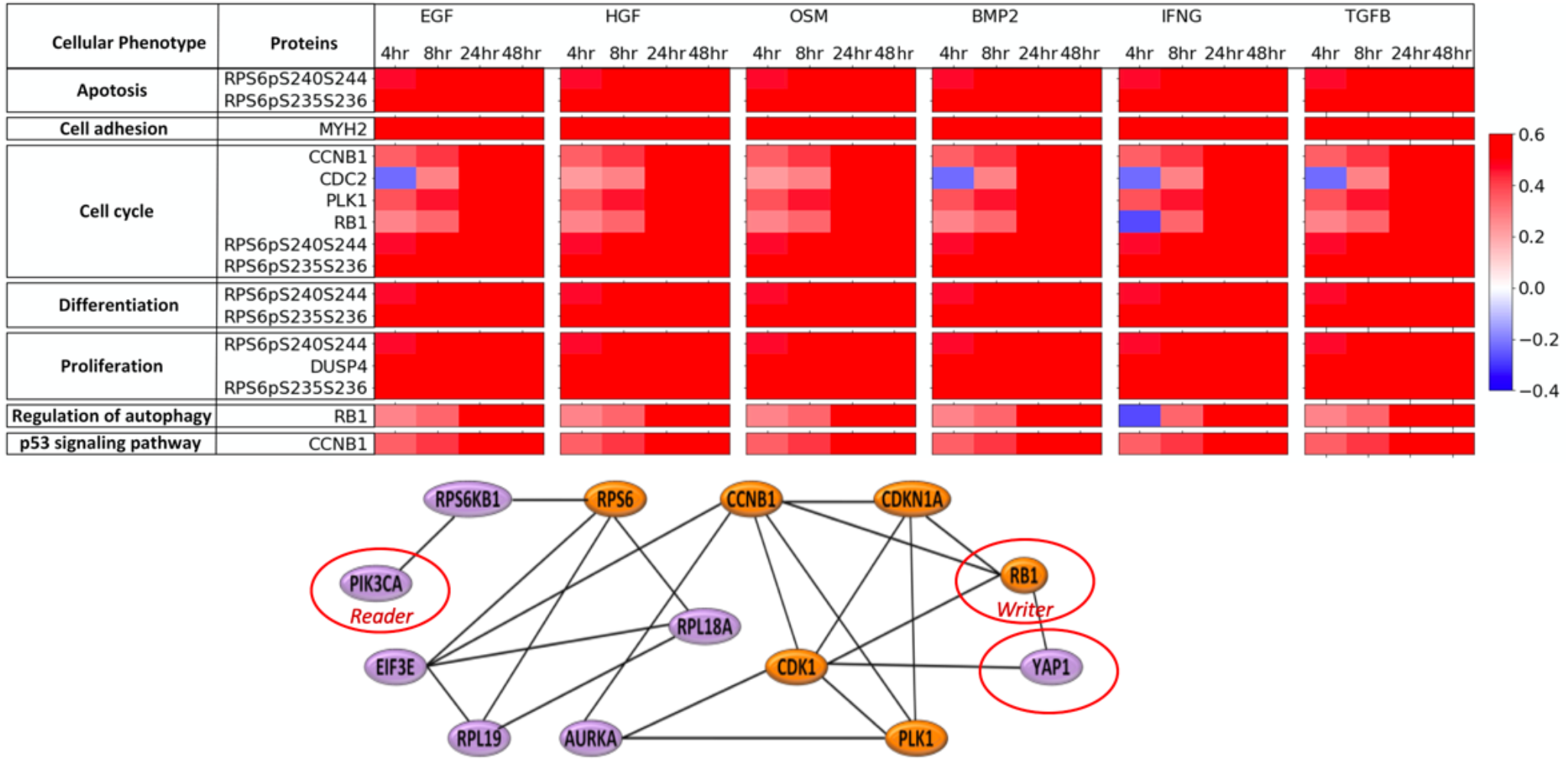
Canonical signaling pathways. Ligands specific signaling pathway profile associated with the enriched proteins at various time points, where positive and negative fold changes are indicated by red and blue colors, respectively. Showing the protein-protein interaction network among five enriched proteins (orange, obtained based on NTF) and seven inferred (purple) proteins.

### Transcriptomic expression analyses show enrichment of cell cycle pathway and corroborate with proteomics results

*Differential analysis:* To provide validation for the ligand-protein correlations produced by the HOCMO, we performed differential expression analysis on RNAseq data to observe the changes in gene expression caused by the growth ligands. We observed CCNB1, CDC2 (also known as CDK1), PLK1, and DUSP4 were significantly (p < 0.05) upregulated across all ligands at 24 hours (Figure 7A). We displayed the expression coverage of each ligand at the loci of these genes, where a higher coverage reflects greater gene expression (Figure 7B). We see that coverage is noticeably higher in the ligand-treated samples compared to PBS control. The upregulation of these genes corroborates with the HOCMO results, which found the proteins encoded by these genes to be correlated with each of the ligands. However, further work must be done in order to completely validate the results of the tensor.

**Figure 7:**
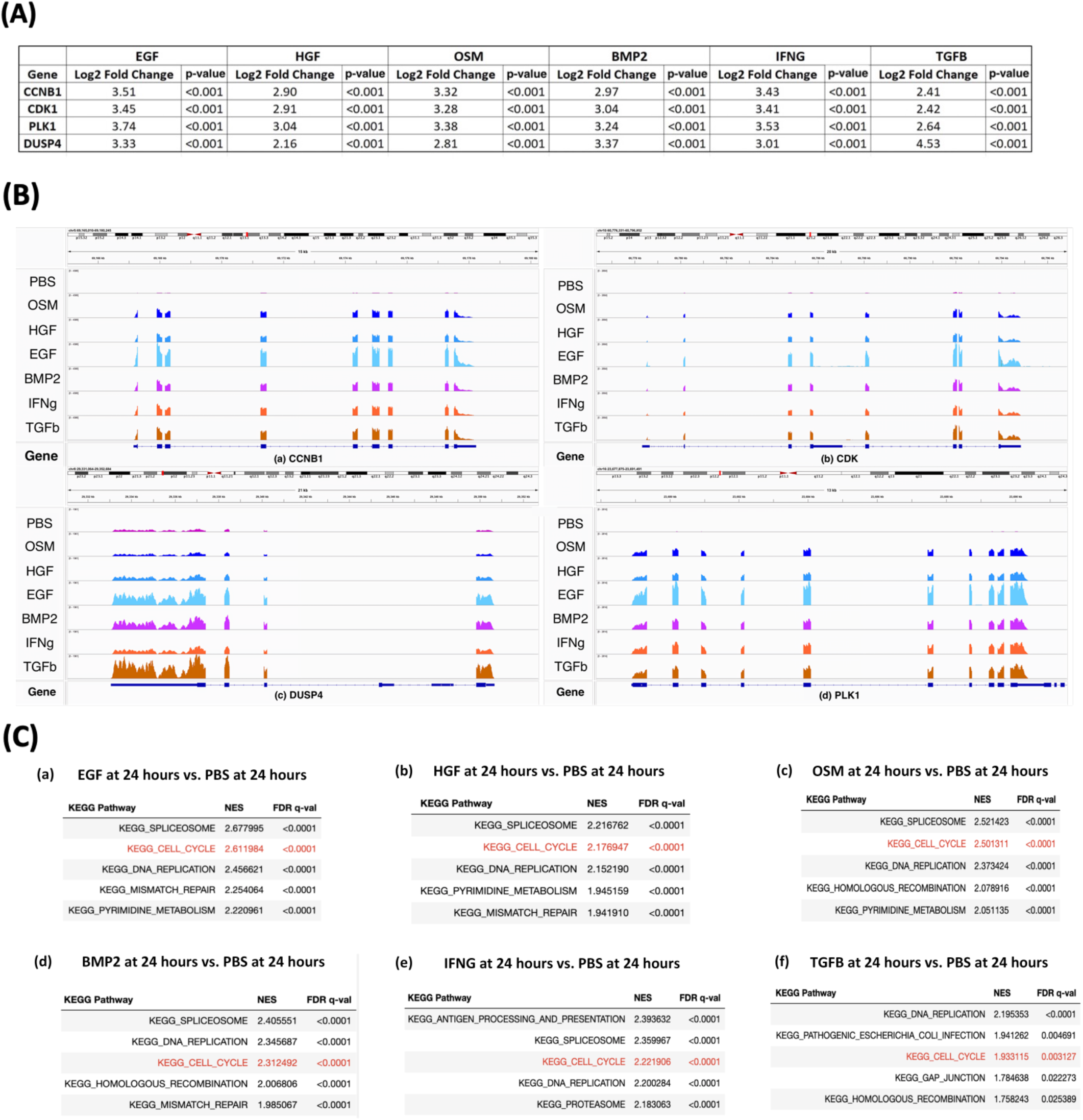
Transcriptomic analyses. (A) Table showing differential expression analysis results of identified enriched (statistically significant, p-value < 0.05) genes treated with ligands at 24 hours. (B) Differential expression analysis. Coverage histograms for each ligand and control at 24 hours showing loci of (a) CCNB1, (b) CDK1, (c) DUSP4, and (d) PLK1. Control is shown on the top track, with the ligands below. Peak height corresponds with expression levels. (C) Top signaling pathways associated with these enriched genes using KEGG pathways.

*Gene Set Enrichment Analysis:* We further performed gene set enrichment analysis on the transcriptomic (RNAseq) data to find out which pathways were most affected by each ligand treatment. We identified top 5 enriched KEGG pathways for each ligand at 24 hours (Figure 7C). Many of the same pathways are highly enriched across all ligands, suggesting that the ligands affect the phenotype of breast cells through similar mechanisms. Cell cycle in particular stands out because the signaling proteins from the HOCMO are key players in mitosis and cell proliferation.

## Discussion

An NTF-based model is a new way to represent higher-order correlations among various biomolecular signals in the breast tumor microenvironment. Further, elucidating the regulatory principle governing the higher-order signaling analysis among phosphoproteins and histones may provide insight into the structural basis of ECM and chromatin remodeling, uncovering a paradigm in cancer cell signaling. Higher-order assemblies are an essential aspect of many biological processes because they enable the formation of precisely organized molecular machines from constituents present in inactive states at low concentrations, such as protein abundances, to promote biochemical reactions in cells. We have introduced an efficient new tensor decomposition method, HOCMO, based on proteome scale signals of MS expressions, and on histones’ phosphoprotein-binding relations among proteins, thus using histones to separate proteome scale nondirectional tensors into mathematically defined decorrelated components. We have demonstrated the utility of NTF on three-dimensional higher correlation data to uncover higher-order protein-ligand and histone-ligand sub-tensors, and the tripling relations among them with biological and statistical significance. This approach focuses mainly on identifying higher-order signaling within individual pathways in the cellular system (intra-pathway signaling) and higher-order signaling between these transduction signaling pathways (inter-pathway signaling), as well as on identifying multiple histone signatures. The proposed tensor decomposition is a simple approach to extract the orthogonal decomposition in analyses of higher-order proteins and histones correlations in proteome-scale breast cell line data. These components and tripling among them include reconstruction of higher-order signaling within and between pathways from nondirectional tensors of higher correlations. The evaluation on our NCP tensor reconstruction, time, and factors shows slightly better performance than the baseline TensorLy (Figure 3A). The result shows the possible onset of breast cancer initiation at 24 hours of treatments for all growth ligands (Figure 4C). The result also uncovers new higher-order coordinated differential relations among the effects of specific growth ligands on proteins contributing to specific cellular phenotypes, including Apoptosis, cell adhesion, cell cycle, differentiation, proliferation, regulation of autophagy and p53 signaling pathway (Figure 6).

Our analysis also reveals possible effects of the growth ligands on phosphoprotein mediated histone regulations that may alter chromatin structure. For instance, we observe significant effects of all ligands on enriched phosphoproteins, namely CCNB1, PLK1, RB1, DUSP4 and RPS6 regulating H3K9me2S10ph1K14ac0 and H3K9me3S10ph1K14ac0 marks to closed conformation (heterochromatin status) starting at 24-hour (Figure 5b). We also observe all ligands on these same phosphoproteins regulating the H4(20to23)K20me0 mark to an open conformation (Figure 5d). From protein-protein interaction analyses (Figure 5f), we further infer that the YAP1 protein, a Hippo signaling coactivator and a major regulator of cancer stemness binds directly with RB1 (regulator involves in heterochromatin formation), indirectly with PIK3CA (chromatin remodeler), by directly interacting with RPS6 (ribosomal protein), suggesting its possible role in chromatin remodeling affecting the cell proliferation, regulation of autophagy, and differentiation in breast cancer when treated with all the ligands (Figure 5b, c, d).

Through the differential gene expression analysis, we further corroborated with the correlations found between ligands and proteins by the NTF-based model. Namely, genes CCNB1, CDK1, PLK1, and DUSP4 were significantly upregulated across all 6 ligands at 24 hours, and the model predicted that all 6 ligands have a significant effect on the proteins encoded by these genes. Similar gene expression profiles have also been reported in other studies, providing further support for the reliability of the model^32–36^. However, for robust support for the model to be achieved we must analyze additional datasets from the LINCS study and conduct validation experiments. Dysregulation of cell cycle genes is a hallmark of tumor cells across various cancer types including breast cancer, and cyclins and cyclin-dependent kinases in particular are important markers. High CCNB1 expression is associated with poor survival in breast cancer^37^, and cyclin expression levels are even used to subtype breast cancers^38^. While much is known about their expression profiles and how they interact with other proteins in breast tumor cells, their involvement in histone modification and downstream epigenetic effects on the tumor microenvironment are relatively unknown. The protein-histone correlations produced by the NTF-based model suggest that these genes have some sort of interaction with the histone modifications H3K9me2S10ph1K14ac0, H3K9me3S10ph1K14ac0, H4(20to23)K20me0, H3K18ub1K23ac0, and H3K4ac1. Based on current knowledge, there are a few possible mechanisms by which cell cycle genes may be involved in histone modification. CDK1 and PLK1 have been found to affect expression of histone methyltransferases via phosphorylation^39–41^. The presence or absence of methyl groups on histones can alter the conformation of chromatin, causing downstream regulatory effects on many genes and drastically altering the tumor microenvironment. The genes may be indirectly modifying histones, like in the above-mentioned studies, or it may be direct. In order to elucidate the details of protein-histone correlations, we plan to use the ATACseq data from the LINCS study to understand the chromatin accessibility landscape of the cells, and perform gene knockout experiments with CUT&RUN^42^ to find the direct effects of these genes on the histones from the model. Ultimately, we will be able to create a network of these genes, their transcription factors, and the histones.

## Supporting information

Supplemental files for tables

## Code Availability

Code is available on GitHub at https://github.com/smollahlab/HOCMO. This includes the code for the HOCMO package available on PyPi, as well as the analysis of RPPA and GCP LINCS data using HOCMO.

## Acknowledgments

This work was supported by Washington University in St. Louis School of Medicine Genetics departmental fund. We want to thank NIH LINCS consortium for making these data available, and Min Shi for helping with figure generation.

## Author Contributions

S.M. conceived and designed the project and came up with the conception. C.L. K.L, and S.M implemented HOCMO. R.S. performed transcriptomic analysis. S.K. performed GSEA analysis. R.S, S.M. performed data interpretation. S.M. wrote the manuscript and supervised the research.

